# A 3D High Resolution Generative Deep-learning Network for Fluorescence Microscopy Image

**DOI:** 10.1101/743179

**Authors:** Zhou Hang, Li Shiwei, Huang Qing, Liu Shijie, Quan Tingwei, Ruiyao Cai, Ali Ertürk, Zeng Shaoqun

## Abstract

Deep learning technology enables us acquire high resolution image from low resolution image in biological imaging free from sophisticated optical hardware. However, current methods require a huge number of the precisely registered low-resolution (LR) and high-resolution (HR) volume image pairs. This requirement is challengeable for biological volume imaging. Here, we proposed 3D deep learning network based on dual generative adversarial network (dual-GAN) framework for recovering HR volume images from LR volume images. Our network avoids learning the direct mappings from the LR and HR volume image pairs, which need precisely image registration process. And the cycle consistent network makes the predicted HR volume image faithful to its corresponding LR volume image. The proposed method achieves the recovery of 20x/1.0 NA volume images from 5x/0.16 NA volume images collected by light-sheet microscopy. In essence our method is suitable for the other imaging modalities.

## Introduction

High resolution volume imaging occupies important position on biomedical imaging field, with the sake of providing more detailed information of the specimens. In the field of optical technique and molecule labeling, some microscopy methods have achieved an unprecedented progress, enabling us to observe thick tissues and whole organs at submicron resolution [1–3]. For example, the morphology of a single neuron across the whole brain [4] or even of the nerve system throughout the entire mouse body can be imaged at the cellular resolution [5, 6]. Limited by Lagrange invariant [7], it is hard to improve both field of view and resolution in optical system. Thus when reaching an imaging depth over working distance of optical lens, optical system often works at low resolution level. This dilemma obstructs the development of high resolution large volume imaging. Common ways to optimize both field of view and resolution, depend on the complicated optical setups, precise control in sample preparation and in imaging process [8]. The generation of high-resolution datasets in large-volume imaging is still not easy to reach.

In recent years, image processing techniques based on deep learning have been successfully applied in optical microscopy imaging [9–12]. Among them, image super resolution technique uses a deep neuronal network to describe the complicated connections between the registered pair of LR and HR images without precise physical model of imaging process. It achieves the transformation of 2D diffraction limited images into the super resolution ones or even in cross-modality [13], however, meets challenge in volume images. A bottleneck for training super resolution network is lacking of massive precisely registered LR and HR volume image pairs. Two following cases deteriorate precise volume image registration. One is that some detailed features in HR volume images cannot be found in LR volume images, the other is that there are the dynamically changes for the labeled fluorescence in the sample and the point spread function of the imaging system. To solve this problem, one way is using some imaging degrading models to generate LR images from experimental HR images [14, 15]. The image degrading model still cannot describe the nonlinear, complicated, and various mapping between LR and HR images, which may cause some poor recovery of HR images in some cases, especially for large HR volume images.

Here, we present a new three-dimensional (3D) high resolution image generation method based on dual-GAN [16], called 3D-dualHRGAN, to recover HR volume images from LR volume Images. The proposed method is model-free and does not require precise image registration in constructing training datasets. One part of 3D-dualHRGAN is traditional GAN to generate high-resolution-like images, avoiding direct comparison of LR and HR images pairs. Furthermore, our network includes the cascade of super-resolution and image degrading network, which makes the contents of generated HR images consistent with that of LR images. For our network, precise image registration is not necessary. We applied our deep neuronal network on neuronal images collected by light sheet microscopy. We demonstrated that 20x volume images are successfully recovered from 5x volume images. The results indicate that one voxel information in LR volume images can be successfully assigned into 3×3×3 voxels in HR volume. This ratio, 27:1, is far more than 4:1 for other super resolution method [15].

## Methods

For this study, we used an adult Thy1-GFPM mouse transgenically expressing green fluorescent protein (GFP) in a population of neurons [17]. In particular, the GFP expressed in the animal was boosted and the animal was cleared using the whole body vDISCO pipeline [6]. We subsequently imaged the same region of the brain of the animal at the level of the cortex and corpus callosum by using two light-sheet microscopes to achieve the acquisition of LR and HR images respectively. The Zeiss Light-sheet Z.1 microscope was used to obtain the low resolution scans, while the LaVision BioTec Light-sheet Ultramicroscope II was employed for the acquisition of high resolutions cans. Detailed data collection information can be found in the Supplementary.

After using the method described above to generate the LR and HR volume neuronal images, we manually ascertained some feature points in the thick neurites and somas that can be found in both LR and HR volume image, and got the feature point pairs. The feature point pair includes the two positions of feature points in LR and HR volumes. According to the feature point pairs, we calculated their transforming matrix. Using the transforming matrix, we achieved the registration from LR images and HR images. This registration is coarse, and the registration error is about dozens of voxel’s size. We divided the coarsely registered LR and HR volume images into a series of sub-blocks and used these sub-blocks to construct training dataset. Considering that 3D convolution layers occupy lots of GPU resource, The size of LR and HR samples are set as 32×32×32 and 96×96×96 respectively.

3D-dualHRGAN includes four convolutional neuronal networks: HR image generator (HR_G), image degrading generator (LR_G), HR image discriminator (HR_D) and LR image discriminator (LR_D), as shown in Fig. 1(a). HR image generator and discriminator constitute the primal GAN in which discriminator HR_D discriminates between the outputs of generator HR_G and the real HR images. LR image is transformed to predicted HR image by HR image generator. By optimizing discriminator HR_D, predicted HR images are gradually well fit the real HR images. Similarly, the dual GAN includes degraded image generator and LR image discriminator, and make predicted LR images, i.e., output of generator LR_G, close to the real LR images. The generators HR_G and LR_G are closely related. When a LR image inputs the cascade of LR_G and HR_G, called cycle-consistent network, (Fig. 1(b)), the ideal case is that this LR image is equal to its output. Similarly, for the other cycle-consistent network, i.e., the cascade of HR_G and LR_G, its output(Fig. 1(b)) should also be consistent with its input.

**Fig. 1.**
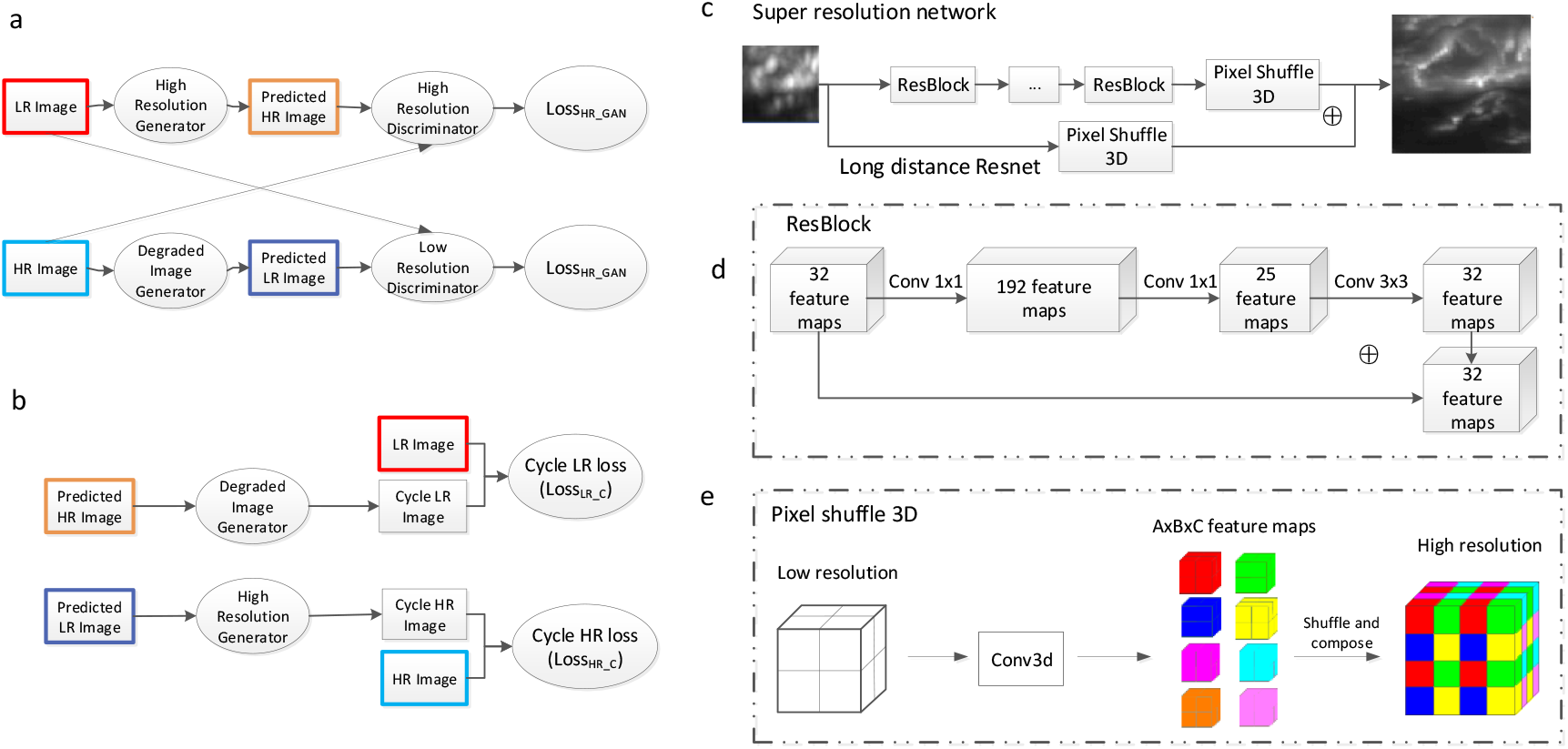
The structure of 3D-dualHRGAN. (a) High/Low resolution generator translates low/high resolution to predicted high/low resolution image. Real high/low resolution image and predicted high/low resolution images are input to high/low resolution discriminator. (b) High resolution/degraded image generator translates predicted low/high resolution images to cycle low/high resolution images. Cycle losses (L1 loss) are applied to keep consistency between cycle low/high resolution images and corresponding real low/high resolution images. (c) High resolution generator includes 8-layer Resblock and a long distance Resnet structure. The last layer is pixel shuffle 3D. (d) The shape of Resblock is hourglass which is called linear low-rank convolution. (e) Pixel shuffle 3D are applied as up sample layer. It unpacks low resolution feature maps into pixels and fills them in high resolution image equidistantly.

Considering the characteristics of 3D-dualHRGAN, as described above there are four loss functions used to train 3D-dualHRGAN. The two loss functions used in discriminator HR_D and LR_D are defined as

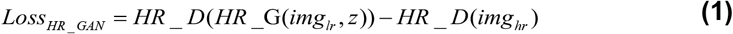

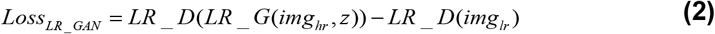

*img_lr_* and *img_hr_* are low-resolution and high-resolution volume images respectively, and z is random noise. The format of these two losses is advocated by WGAN training strategy [18, 19]. Compared with sigmoid cross-entropy loss used in traditional GAN, this format has faster convergent speed and better translation results. The other two loss functions with generator HR_C and LR_C are defined as

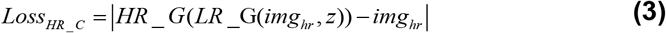

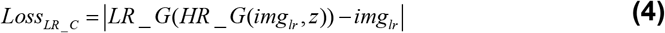

| | represents L1 distance. The design of these two losses is originated from image super-resolution model [18]. By minimizing these two losses, the output of generator HR_G (predicted HR volume image) can be well consistent with its input (LR volume image), and the artifacts in predicted HR volume image has been vastly suppressed. These four loss functions are combined together for the total object that trains 3D-dualHRGAN, given below

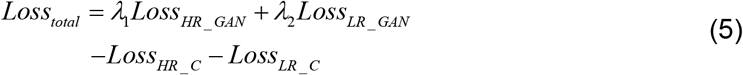

Here, *λ*_1_ and *λ*_2_ are two hyper parameters experimentally determined to balance the four losses. In our analysis, they are set to be 10 which make the generator loss keep the same level with the discriminator.

We designed the generators and discriminators in 3D-dualHRGAN which are suitable for volume image super-resolution.

The HR image generator includes 8-layer short ResBlock and long distance ResNet structure (Fig. 1(c)). Long distance Resnet takes low-level feature information to the last layer and output the final predicted HR images. 8-layer short ResBlock are applied to extract low and high level image feature information. As shown in Fig. 1(d), the Resblock has hourglass-shape that has been widely used in 2D image super-resolution analysis, which is known as linear low-rank convolution [20]. In this structure, at first the convolution operations is performed with 192 filters, then it is reduced to 25 filters to shrink the image features. This structure can reach a good high resolution prediction effect with less parameters cost. We adopted weight normalization layer rather than batch normalization to make the network more flexible. In the up-sampling layer, we adopted voxel shuffle procedure to integrate *A*×*B*×*C* feature maps (of which the size is *W*×*H*×*D*) into high resolution image, of which size is (*A*×*W*)×(*B*×*H*)×(*C*×*D*). Each feature map is unpacked into pixels and filled in high resolution image equidistantly like traditional pixel shuffle [21], as shown in Fig. 1(e).

The image degrading generator is used to map a HR volume image to a LR volume image. It is a convolutional neuronal network, consisting of 5-layer 3×3×3 convolutional layers. The stride of last convolution layer is set as 3. This layer is used as down-sample layer. The discriminator HR_D and LR_D are also convolutional neuronal networks. Discriminator HR_D includes 11 convolutional layers and 5 down-sampling layers. The discriminator LR_D includes 5 convolutional layers and 3 down-sampling layers. We adopted WGAN-GP method [18, 19] to train GAN networks. This requires that these two discriminators directly output patches. So, in these two discriminators, we deleted the sigmoid layer where the activate functions are set.

Our 3D-dualHRGAN transforms 5x volume images to 20x volume images. By this transformation, in the x-axis, y-axis and z-axis, the voxel size of HR volume image increases to 3, 3, and 3 times, namely, the total number of voxels of LR and HR volume images has the ratio of 27:1. Therefore, when the size of LR and HR volume images is large in the training set, huge computational resource is required. Considering this point, we set the size of LR and HR volume images in training set to 32×32×32 and 96×96×96 respectively, as previously illustrated.

3D-dualHRGAN consists of two GAN networks and two cycle-consistent networks. We used WGAN-GP [18, 19] method to train the two GAN networks. The corresponding algorithm is RMSprop, and the learning rate starts with 10^−4^. The classical training method with Adam algorithm is used for the two cycle-consistent networks. The start learning rate ranges from 10^−4^ to 10^−3^. In the training procedure, we fix the generator HR_G and LR_G to optimize the discriminator HR_D and HL_D by minimizing the object function (1-2) and then, optimize the generator HR_G and LR_G in the condition of fixing these two optimized discriminators. In meantime, the gradient penalty process is applied to keep training process stable. Repeating this procedure makes the generators and discriminators iteratively optimized until the total loss keeps stable.

## Result

We demonstrated that the proposed method can improve the volume image resolution. We used a set of registered volume images which include soma and sufficient neurites for this demonstration. LR volume images were acquired with 5x/0.16NA objective, and had the same size of 100×100×100. Their corresponding HR volume images (same anatomic region) were acquired with 20X/1.0 NA objective, and considered as the ground truth. The maximum intensity of XY, XZ, YZ projections from the LR, HR and predicted HR volume images have been shown in Fig. 2. We observed that the volume image resolution, especially axial resolution was very low, which caused some neurites not being identified. The predicted HR volume images (the second column in Fig. 2(a)), as the outputs of 3D-dualHRGAN, had the size of 300×300×300, 27 times the total number of voxels in the input images. In the predicted images, the spatial resolution was substantially improved, and more details were visible. To furthermore illustrate the resolution improvement, we compared the predicted images to the real HR images. The image intensities on the color lines were extracted from the inputs, from the predicted and from the real HR images, and all have been plotted in the fourth column of Fig. 2(a). The peak widths of image intensity curves from the predicted image were smaller than LR images and close to HR images. We noticed there were signal loss and position shift in LR and predicted HR images. It was due to the lower imaging ability for LR images and registration bias. From XY, XZ and YZ projections, we measured 20 neurite peak widths from different neurites of 4 sets of images. The values of 60 peak widths have been sorted and normalized. Then the preprocessed datasets were plotted on Fig. 2(b). Blue, red green curves were corresponding to LR images, HR images and predicted HR images. The neurite peak widths of HR and predicted HR images were closer than LR and HR images. The plotted curves in Fig. 2(b) shows the PSF reduction feature of predicted HR images produced from our network, especially on XZ and YZ projection.

**Fig. 2.**
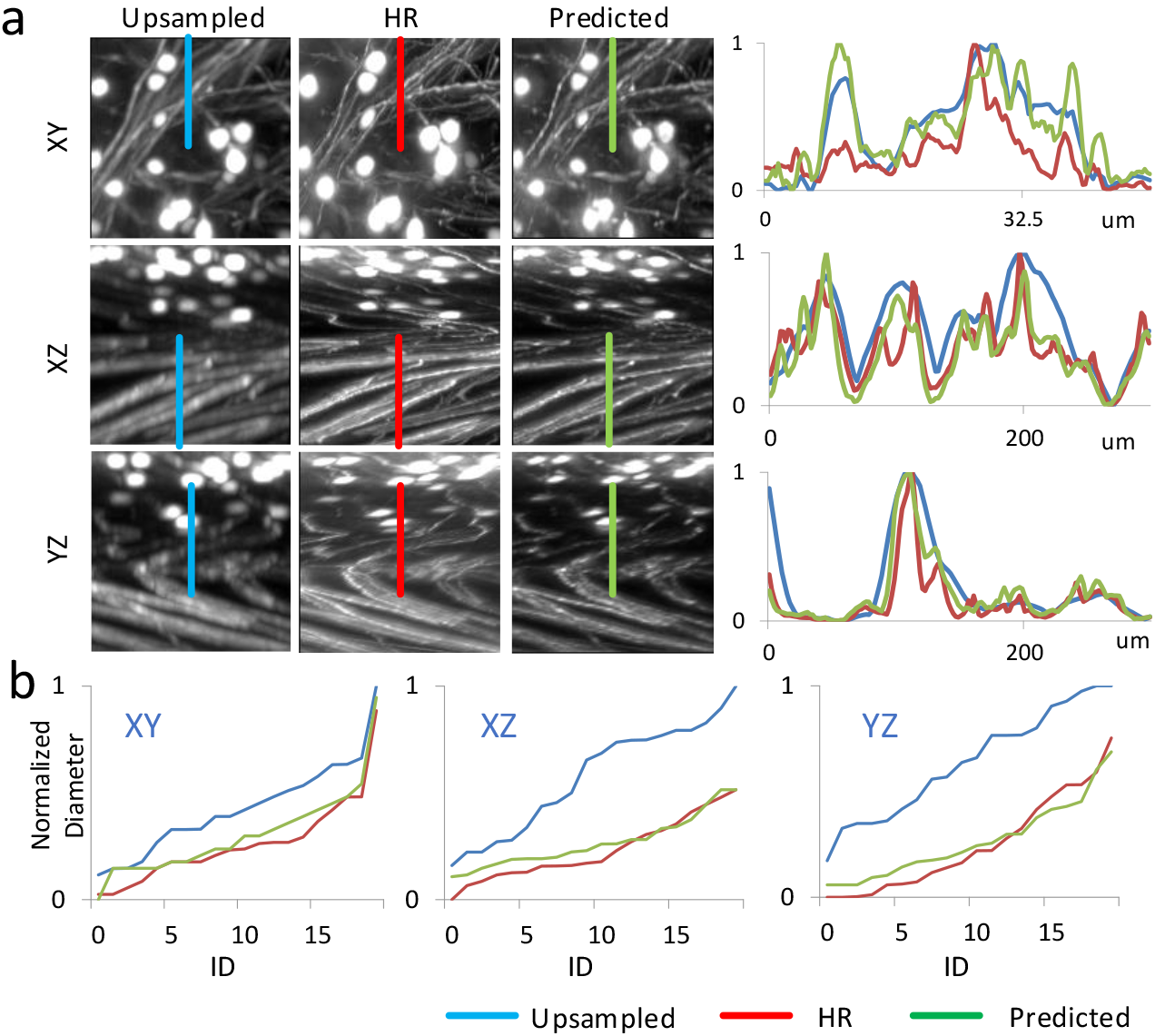
(a) Comparison of bilinear up-sampled images, predicted images and high resolution images. The fourth column shows the signal distribution on color lines for corresponding images. (b) we measured 20 peak widths on XY, XZ YZ projection from different neurites. The measured data are normalized, sorted and then plotted. In (a) and (b), blue, red, green curves are corresponding to up-sampled images, predicted images and high resolution images.

We applied the proposed method to improve the resolution of large-volume neuronal images. The dataset comes from a brain region showing cortex and corpus callosum and has the size of 660×660×160. Its predicted HR image has the size of 1980×1980×480. We compared the input and the output of the proposed network (Fig. 3). The neurites become thin and some touched cells are apart after the prediction (yellow arrows). There are almost no artifacts in the predicted HR images. In addition, the weak signals in the LR volume image can also be successfully predicted in its predicted HR volume images (pink arrows). These results can also be illustrated in their XZ cross-sections(Fig. 3(b)). From the presented results, we conclude that the predicted HR images not only strictly originate from the LR images but also the resolution of the LR images was substantially improved.

**Fig. 3.**
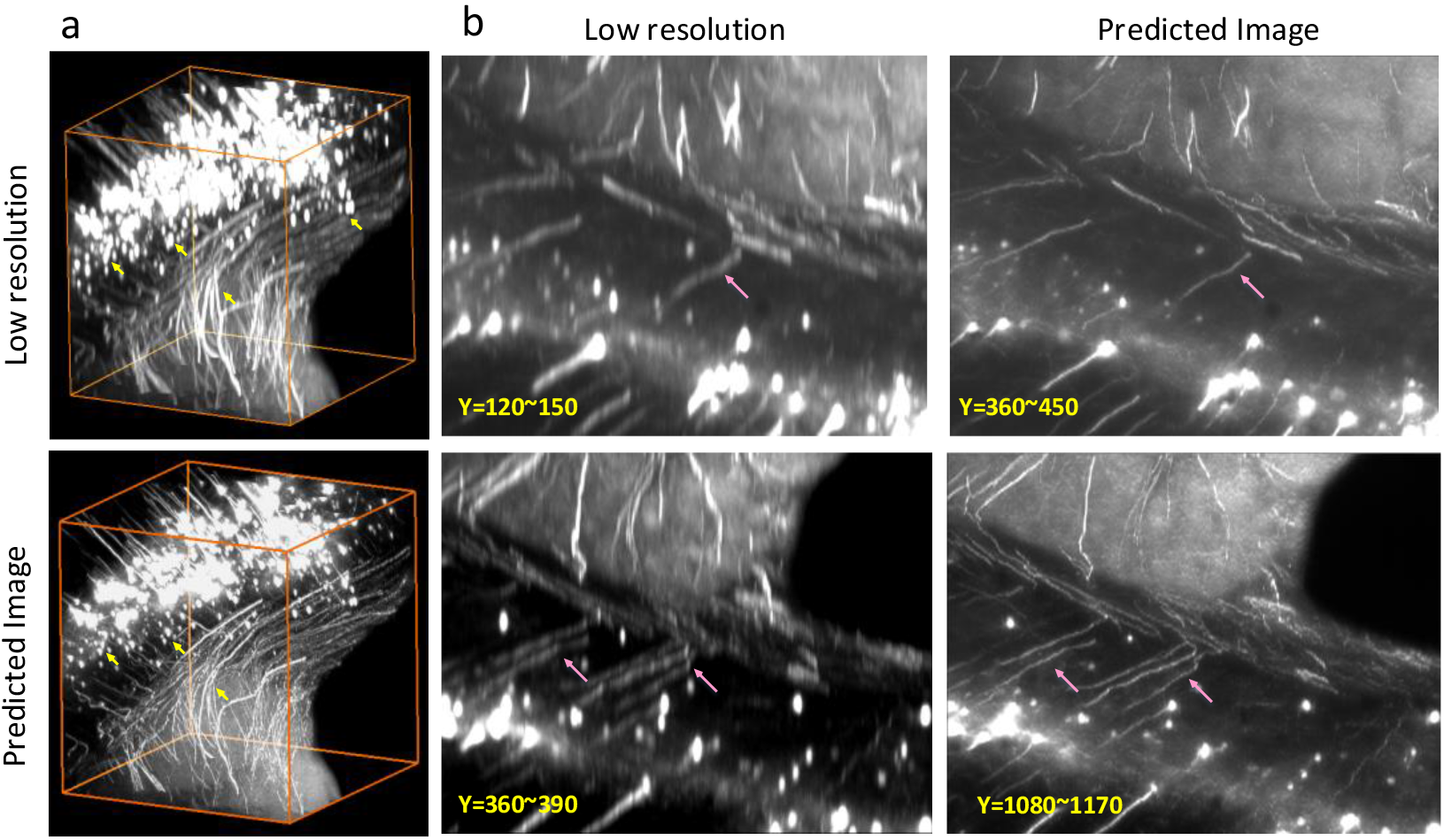
(a) 3D LR large image (up) and corresponding predicted HR image (down). The size of LR image is 660×660×160. The size of predicted HR image is 1980×1980×480. In order to have a more detailed observation, the Z axis of the volume rendering is scaled up to 4 times. Yellow arrows in LR images point out the areas where soma and neurites touched. In predicted HR image, yellow arrows point out corresponding areas where soma and neurites are apart away. (b) shows the maximum intensity projection images of 3D LR image (left column) and corresponding predicted HR image (right column) on XZ plane. Projection images in (b) are corresponding to XZ plane at different Y value. Pink arrows show that weak signal areas can be predicted well using our method.

In conclusion, we present a new 3D high resolution generative Deep-learning Network called 3D-dualHRGAN for recovering HR volume images from LR volume images. This method introduces dual-GAN network to avoid precise registration process of HR and LR volume image pairs in standard deep neuronal networks, and meanwhile captures well the mappings from LR to HR volume images. The proposed network can make the predicted HR volume images consistency with corresponding LR volume images. We applied the proposed network to neuronal images, and the trusted HR volume images can be obtained. As this method does not rely on prior knowledge of imaging modality, this method is hoped to be applied in other biological volume imaging systems.

## Funding

National Program on Key Basic Research Project of China. (Grant No. 2015CB7556003) and Director Fund of WNLO. Vascular Dementia Research Foundation, Synergy Excellence Cluster Munich (SyNergy to A.E.), ERA-Net Neuron (01EW1501A to A.E.), Fritz Thyssen Stiftung (A.E., Ref. 10.17.1.019MN), DFG (A.E., Ref. ER 810/2-1 and TRR127 to E.W. and E.K.), NIH (A.E.), and Helmholtz ICEMED Alliance(A.E.).

## Acknowledgment

We thank Maximilian Gorelashvili from Carl Zeiss Microscopy GmbH for the help in obtaining the LR scans.

