## Supplementary for "A 3D High Resolution Generative Deep-learning Network for Fluorescence Microscopy Image"

The biological samples used in this study are coming from a Thy1-GFPM mouse of 2-3 months old, expressing green fluorescent protein (GFP) in a subset of neurons (mostly somatosensory and motor neurons) [1]. The animal experiments were conducted according to institutional guidelines: Klinikum der Universität München / Ludwig Maximilian University of Munich and after approval of the Ethical Review Board of the Government of Upper Bavaria (Regierung von Oberbayern, Munich, Germany). After inducing deep anesthesia using midazolam-medetomidine-fentanyl (1ml/100g of body mass; i.p.), the mouse was intracardially perfused with room temperature 0.1 M PBS containing heparin (10 U/ml of Heparin, Ratiopharm; ~110 mmHg pressure using a Leica Perfusion One system) for 5-10 minutes to wash out the blood and this was followed by perfusion with 4% paraformaldehyde (PFA) in 0.1 M PBS (pH 7.4) (Morphisto, 11762.01000) for 10-20 minutes. Next the skin was removed from the body of the animal and the skull was opened on the dorsal side to expose the dorsal cortex of the brain. Then the body was post-fixed in 4% PFA for 1 day at 4 °C and subsequently washed with 0.1 M PBS for 10 minutes 3 times at room temperature.

The transgenically expressed GFP of the animal was boosted and the animal was cleared by the whole body vDISCO method [2] using 35µL of Atto647N conjugated anti-GFP nanobooster (Chromotek, gba647n-100). In particular, we used 20-35 µg of the nanobooster in 250 ml (0.08-0.14 µg/ml), 1:7000 in dilution, (stock concentration 0.5 – 1 mg/ml). Then, after clearing the brain was dissected out from the animal for imaging.

The imaging of the brain was done using two microscopes for the LR and HR respectively.

For LR images, we used Zeiss Light-sheet Z.1 microscope with the commercially available specifications to obtain the LR scans. In particular, the objective used was Zeiss Z.1 detection optic 5x/0.16 NA (water, clearing n=1.45). The zoom was 1x

For HR images, we used LaVision BioTec Light-sheet Ultramicroscope II with the commercially available specifications to obtain HR scans. In particular, the objective used is Zeiss 20x Clr Plan-Neofluar/0.1 NA [WD 5.6 = mm]). The zoom was 1x.

1. G. Feng, R. H. Mellor, M. Bernstein, C. Keller-Peck, Q. T. Nguyen, M. Wallace, J. M. Nerbonne, J. W. Lichtman, and J. R. Sanes, Neuron **28**, 41-51 (2000).

2. R. Cai, C. Pan, A. Ghasemigharagoz, M. I. Todorov, B. Forstera, S. Zhao, H. S. Bhatia, A. Parra-Damas, L. Mrowka, D. Theodorou, M. Rempfler, A. L. R. Xavier, B. T. Kress, C. Benakis, H. Steinke, S. Liebscher, I. Bechmann, A. Liesz, B. Menze, M. Kerschensteiner, M. Nedergaard, and A. Erturk, Nat Neurosci **22**, 317-327 (2019).
